# A Data-Driven Evaluation of the Size and Content of Expanded Carrier Screening Panels

**DOI:** 10.1101/430546

**Authors:** Rotem Ben-Shachar, Svenson MS Ashley, James D. Goldberg, Dale Muzzey

**Author notes:** Correspondence: Dale Muzzey, 1-888-268-6795.

## Abstract

**Purpose:** The American College of Obstetricians and Gynecologists (ACOG) proposed seven criteria for expanded carrier screening (ECS) panel design. To ensure that screening for a condition is sufficiently sensitive to identify carriers and reduce residual risk of non-carriers, one criterion requires a per-condition carrier rate greater than 1-in-100. However, it is unestablished whether this threshold corresponds with a loss in clinical detection. The impact of the proposed panel-design criteria on at-risk couple detection warrants data-driven evaluation.

**Methods:** Carrier rates and at-risk couple rates were calculated in 56,281 patients who underwent a 176-condition ECS and evaluated for panels satisfying various criteria. Condition-specific clinical detection rate was estimated via simulation.

**Results:** Different interpretations of the 1-in-100 criterion have variable impact: a compliant panel would include between 3 and 38 conditions, identify 11%-81% fewer at-risk couples, and detect 36%-79% fewer carriers than a 176-condition panel. If the carrier-rate threshold must be exceeded in all ethnicities, ECS panels would lack prevalent conditions like cystic fibrosis. Simulations suggest that clinical detection rate remains >84% for conditions with carrier rates as low as 1-in-1000.

**Conclusions:** The 1-in-100 criterion limits at-risk couple detection and should be reconsidered.

## INTRODUCTION

Carrier screening facilitates reproductive decision-making by identifying couples at-risk for conceptuses affected with autosomal recessive (AR) and X-linked conditions.^1^ Advances in genomic technology coupled with decreasing sequencing costs has led to the advent and adoption of expanded carrier screening (ECS) for tens to hundreds of recessive conditions simultaneously.^2^ For AR conditions, a couple is at risk if both individuals are carriers of the same condition, with the conceptus having a 25% probability of being affected with the condition. For X-linked conditions, a couple is at risk if the female is a carrier: the probability of a male conceptus being affected with the condition can be up to 50%.^3^ The American College of Obstetricians and Gynecologists (ACOG) recently published a committee opinion stating that ECS is an acceptable strategy for carrier screening.^1^ However, no consensus exists on which conditions should be included on an ECS panel.

Rather than prescribing specific conditions for an ECS panel, professional societies have provided general guidelines for ECS-panel design, stressing that panels should maximize clinical utility and not simply follow the paradigm of “the more conditions, the better.”^1,4,5^ Yet there remains no specific recommended set of ECS conditions in part because there are over 1,000 possible single-gene recessive conditions that could be included on a panel,^6^ and it is difficult to determine unambiguous criteria.^7^ As a result, the size and content of ECS panels vary widely across laboratories.^7,8^ In a recent study comparing 16 commercially available ECS offerings, panel size ranged from 41 to 1,792 conditions, with only three conditions screened by all panels.^7^

Discrepancies in panel size and content have led to a growing desire for guidelines that delineate which conditions should be included on ECS panels.^1,8^ Because ECS is commonly performed sequentially (i.e., ECS on initial patient and single-gene screening as needed on the partner), inclusion of each additional condition on a panel poses a trade-off: identification of at-risk couples provides actionable information to guide pregnancy management,^9^ while concomitant detection of carriers can increase patient anxiety,^2,10,11^ require additional counseling,^2,10^ and introduce logistical burden for the provider.^2,11,12^ For many conditions, the benefit of identifying at-risk couples offsets these challenges, but it is unclear when a condition is too rare to warrant screening. To that end, ACOG^1^ and a clinical assessment of ECS panel design^8^ recommend a 1-in-100 carrier frequency threshold for condition inclusion. However, these studies do not quantitatively consider how the 1-in-100 carrier-frequency threshold affects the trade-off between carrier identification and at-risk couple identification.

Professional societies further emphasize that in order for an ECS panel to have high clinical utility, conditions on ECS panels should minimize residual risk (i.e., the risk that a patient carries a pathogenic variant after screening negative).^1,3,4,8^ Genetics professionals recommend that minimal residual risk be achieved by setting a minimum threshold for screening detection rate,^8^ which depends on two compounding factors: analytical detection rate (the ability to accurately detect variants) and clinical detection rate (the ability to accurately determine if a variant is pathogenic or benign). High analytical detection rates have been demonstrated for most conditions on ECS panels,^8,13^ but clinical detection rates have yet to be systematically evaluated.

Building on the guidance of professional societies, a study proposed and applied panel design criteria to seven commercially available ECS offerings, yielding a panel of 96 conditions.^8^ A complementary approach proposed an algorithm to classify condition severity based on disease characteristics,^14^ and developed a methodology to maximize detection of conceptus disease risk while ensuring accurate variant interpretation, culminating in a 176-condition panel.^15^ The nearly two-fold disparity in panel size resulting from these two approaches, both of which applied principled panel-design criteria, underscores the need for greater clarity and objectivity in guidelines.

Given the importance of ensuring that ECS panels maximize clinical utility, a data-driven approach is needed to evaluate the impact of professional-society condition-inclusion criteria on detection of carriers and at-risk couples. We evaluated the most recent guidelines for ECS panel design from ACOG’s 2017 committee opinion,^1^ which recommended that each ECS condition meet several of seven proposed criteria: (1) Have a well defined phenotype, (2) Have a detrimental effect on quality of life, (3) Cause cognitive or physical impairment, (4) Require surgical or medical intervention, (5) Have an onset early in life, (6) Have prenatal diagnosis available, and/or (7) Have a 1-in-100 or greater carrier frequency. We retrospectively analyzed data from a panethnic cohort of over 50,000 patients screened with a 176-condition ECS panel and evaluated how exclusion of conditions that did not meet criteria impacted detection of carriers and at-risk couples. We show that any interpretation and application of the 1-in-100 carrier-frequency threshold limits detection of at-risk couples, and different interpretations cause at-risk couple detection rates to vary by 11-fold. Instead of a carrier-rate threshold, we propose an alternative measure—estimation of a clinical detection rate—to evaluate when a condition is too rare to be included in an ECS panel by quantifying if there is enough evidence to determine the clinical association between detected variants and disease.

## MATERIALS AND METHODS

### Cohort design

We retrospectively analyzed de-identified data from patients who underwent ECS over a 17-month period using a 176-condition panel (Foresight; Myriad Women’s Health, South San Francisco, CA).^15,16^ We constructed two patient cohorts for different purposes, one that focused on individual estimates representative of the U.S. population and another that focused on couples for empirical analysis.

The first patient cohort, consisting of 56,281 patients and used for the majority of analyses, excluded patients who had a family or personal history of disease or reported consanguinity. To be reflective of the general U.S. population, we used U.S. census data to weigh the respective ethnicity-specific carrier frequencies and at-risk-couple rates in this cohort.

The second cohort included couples that received ECS to allow for calculation of the empirical frequencies of at-risk couples per condition. One analysis included all of the couples that underwent ECS (N=11,536) who were identified as at-risk (N=501; Figure S1), but most analyses excluded couples with a personal or family history of disease (8,736 couples among whom 314 were at-risk).

This study was reviewed and designated as exempt from institutional review board (IRB) oversight (as granted by Western IRB on April 23, 2018). All patients provided informed consent for testing and anonymized research.

### ECS condition classification using ACOG guidelines

For each of the 176 conditions on the ECS panel, we evaluated which of the seven ACOG criteria (enumerated in the Introduction) were met (Table S1). These criteria were evaluated by a certified genetic counselor, using refined classification criteria described in Supplementary Text S1.

Because ACOG did not provide specific information in its committee opinion,^1^ we evaluated the criterion suggesting that a condition have a 1-in-100 or greater carrier frequency in the following ways: (1) A condition has a 1-in-100 or greater carrier frequency in any ethnicity, (2) A condition has a 1-in-100 or greater carrier frequency when ethnicities are weighted based on their U.S.-census frequencies, or (3) A condition has a 1-in-100 or greater carrier frequency in all ethnicities. All ethnicities were self-reported; ethnicities and U.S.-census frequencies are provided in Table S2. Because of different inheritance characteristics, X-linked and AR conditions with the same prevalence can have very different carrier frequencies. Therefore, we considered two carrier-rate thresholds for X-linked conditions: (1) a carrier rate threshold of 1-in-100 and (2) a carrier rate threshold of 1-in-10,000, which resembles a 1-in-100 carrier rate for AR conditions by yielding a prevalence of 1-in-40,000. Descriptions of how carrier frequencies and at-risk couple rates were computed are provided in Supplementary Text S2. Condition-specific carrier frequencies and evaluation of each 1-in-100 carrier frequency definition are provided in Table S3.

### Estimation of clinical detection rate

Clinical detection rate is a function of variant frequencies and the ability to correctly classify identified variants as being pathogenic or benign. Given our large cohort, we assume that the majority of pathogenic variants for each condition have been observed and that their frequencies are empirically determined from our dataset. However, we also presume that some rare pathogenic variants have not been observed in our dataset. We account for these unobserved variants by assuming (1) that the number of unobserved pathogenic variants in our cohort is proportional to the number of observed pathogenic variants and (2) that the frequency of unobserved variants reflects the frequency of the least-common observed pathogenic variant (Supplementary Text S3).

In order to model if variants could be correctly classified as pathogenic, we relied on ACMG guidelines for interpretation of sequence variants that suggest that enrichment of variants in affected cases relative to controls is strong evidence for pathogenicity.^17^ We modeled estimated clinical detection rates assuming that a minimum of three or more expected cases reported in the literature are needed to accurately classify a variant as pathogenic, based on ACMG guidelines.^17^ Given the relative pathogenic variant frequencies and the expected number of reported cases worldwide for each condition, we simulated the number of reported cases for each observed variant, assuming that all unobserved variants will have no reported cases. We define estimated clinical detection rates as the sum of adjusted variant frequencies with three or more simulated reported cases (Supplemental Text S3). We repeated the simulations of case reports 10,000 times for each condition and re-estimated clinical detection rates assuming that 1, 2, or 4 reported cases are needed to determine pathogenicity of variants (Supplementary Text S3). All analyses were performed using Python 2.7.10, Numpy 1.13.1, and Pandas 0.20.3.

## RESULTS

### Quantifying impact of ACOG guidelines

We retrospectively analyzed an ethnically diverse cohort of 56,281 average-risk patients who underwent ECS with a 176-condition panel to determine how proposed ACOG panel design criteria would have impacted the detection rates of at-risk couples and carriers of recessive conditions. We further determined how many observed at-risk couples detected by the 176-condition panel would not have been identified if proposed guidelines were strictly followed.

We evaluated the collective impact of six ACOG criteria unrelated to carrier frequency on a panel’s at-risk couple rate and the panel carrier rate, which we define as the proportion of patients in the cohort who were carriers of at least one condition on the panel. Of the 176 conditions on the ECS panel, 172 met all six criteria, with the remaining four conditions not present in childhood in the majority of affected patients (Figure 1A). Limiting an ECS panel to these 172 conditions would reduce the panel carrier rate, the at-risk couple rate, and the number of empirically identified at-risk couples each by 3% (Figure 1B-D). We compared this subpanel to a panel that meets the ACOG ethnicity-specific guidelines for screening outlined in ACOG committee opinion 691 (denoted “ACOG 691”)^18^ and a panel that includes only cystic fibrosis (CF) and spinal muscular atrophy (SMA), the two conditions for which ACOG specifically recommends routine screening in all ethnicities (denoted “CF/SMA”).^18^ For the ACOG 691 panel, carrier identification would be reduced by 77%, at-risk couple identification would be reduced by 66%, and 258 observed at-risk couples (82%) would not have been identified (Figure 1B-D). For the CF/SMA panel, carrier identification would be reduced by 88%, at-risk couple identification would be reduced by 84%, and 286 observed at-risk couples (91%) would not have been identified (Figure 1B-D).

**FIGURE 1.**
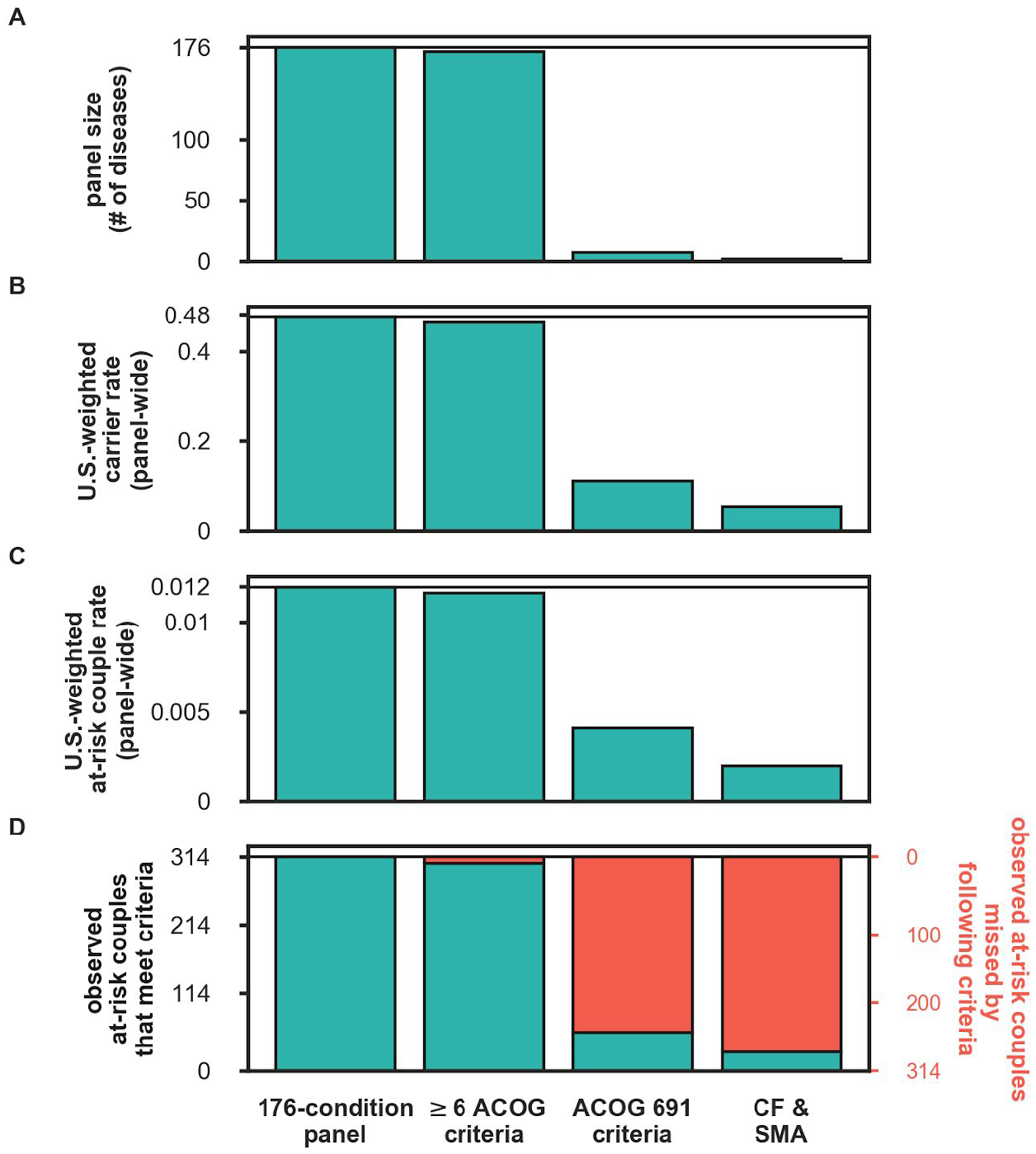
Impact of ACOG guidelines on panel size, carrier rates, and at-risk couple rates. We consider four panels: the full 176-condition panel, the subset of conditions that meet the first six ACOG criteria (excluding the 1-in-100 criteria), the subset of conditions that meet the ethnicity-specific requirements in ACOG guidelines 691,^18^ and a panel that includes only cystic fibrosis (CF) and spinal muscular atrophy (SMA). (A) The number of diseases that meet criteria. (B) U.S.-weighted panel carrier rates. (C) U.S.-weighted at-risk couple rate. (D) The number of observed at-risk couples that would be identified (green) or omitted (red) by the indicated panel. Horizontal lines show respective numbers for the 176-condition panel.

### Large variability in ethnicity-specific carrier rates

To determine the impact of the final ACOG criterion, that ECS conditions have carrier frequencies higher than 1-in-100, we examined the variation in ethnicity-specific carrier rates. We found wide variability in ethnicity-specific carrier rates for five prevalent conditions for which one or more ethnicities have a carrier rate below 1-in-100 (Figure 2A). The relative difference between the maximum and minimum ethnicity-specific carrier rates ranged from 5-fold (*GJB2*-related nonsyndromic hearing loss and deafness) to 52-fold (Hb beta chain-related hemoglobinopathy) (Figure 2B). Even for prevalent conditions such as cystic fibrosis, for which pan-ethnic screening is recommended,^18^ multiple ethnicities have a carrier rate below 1-in-100 (Figure 2A).

**FIGURE 2.**
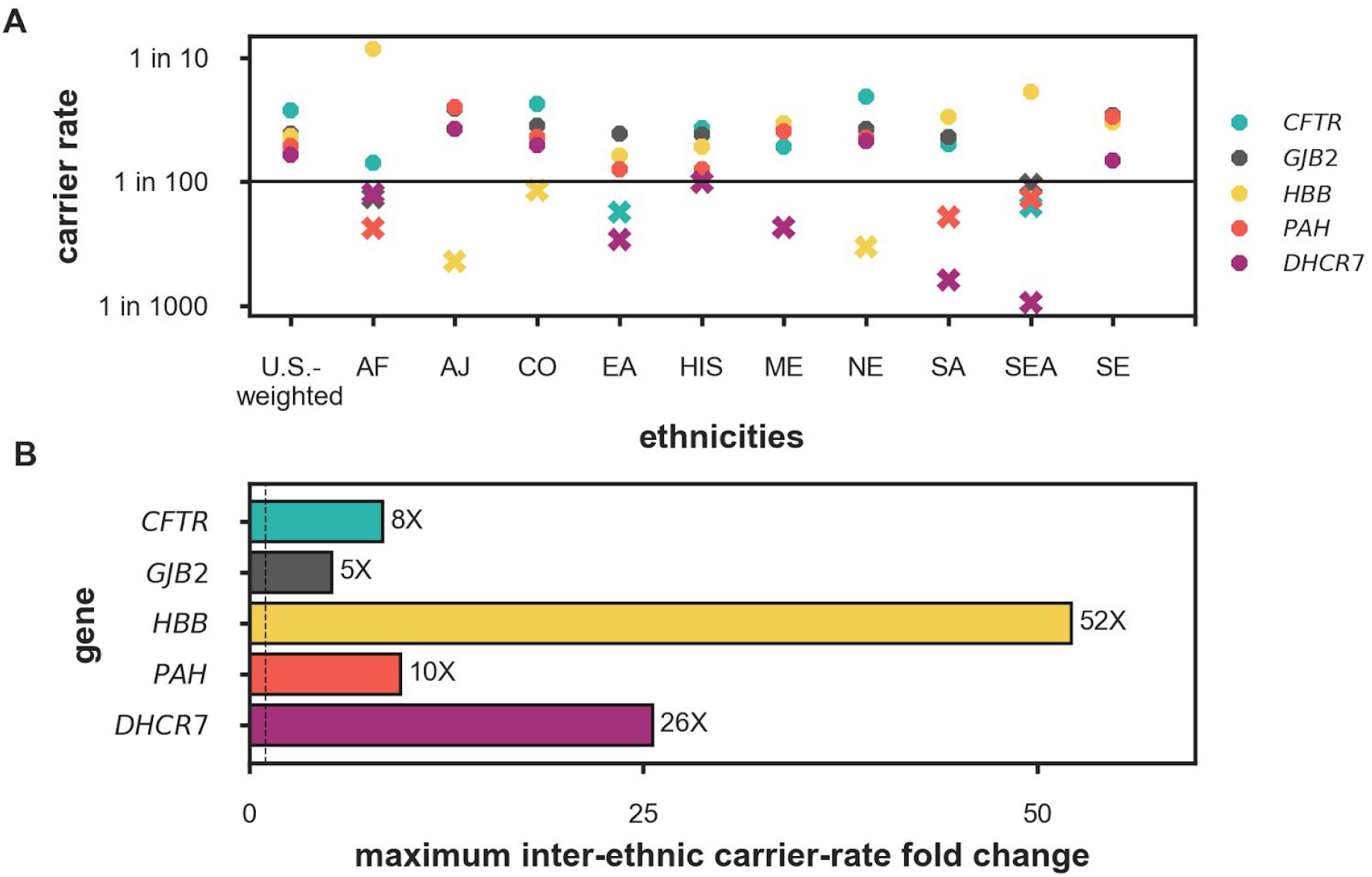
Carrier rates vary widely by ethnicity. (A) Self-reported ethnicity-specific carrier rates for the five conditions with the highest U.S.-weighted carrier rates in the 176-condition panel for which an ethnicity-specific carrier rate is less than 1-in-100 in at least one ethnicity. Ethnicities with carrier rates below a 1-in-100 carrier rate are denoted with Xs; ethnicities with carrier rates above the 1-in-100 carrier rate are denoted with hexagons. (B) The relative difference in carrier rates between the highest and lowest ethnicity-specific carrier rates for each condition in (A). *CFTR*: cystic fibrosis. *GJB2*: *GJB2*-related DFNB1 nonsyndromic hearing loss and deafness. *HBB*: Hb beta chain-related hemoglobinopathy. *PAH*: phenylalanine hydroxylase deficiency, *DHCR7*: Smith-Lemli-Opitz syndrome. AF: African. AJ: Ashkenazi Jewish. CO: Caucasian/Other. EA: East Asian. HIS: Hispanic. ME: Middle Eastern. NE: Northern European. SA: South Asian. SEA: Southeast Asian. SE: Southern European.

### Any interpretation of the 1-in-100 carrier-rate criterion limits detection of at-risk couples

Different interpretations of the 1-in-100 carrier-rate threshold impact identification of carriers and at-risk couples. We quantified the impact of three threshold definitions, ranked from least stringent to most stringent: 1-in-100 carrier rate in any ethnicity, 1-in-100 U.S.-weighted carrier rate, and 1-in-100 carrier rates in all ethnicities (see Methods). We further stratified these three definitions by two carrier-rate thresholds for X-linked conditions: a carrier rate threshold of 1-in-100 and a carrier rate threshold of 1-in-10,000, corresponding to the same prevalence rate as for AR conditions with a 1-in-100 carrier rate. A compliant panel would reduce the panel size from 176 to between 3 and 38 conditions, depending on the definition used (Figure 3A). Carrier rates would be reduced between 36% and 79%, at-risk couple rates would be reduced between 11% and 92%, and between 33 (11%) and 255 (81%) observed at-risk couples would not be identified (Figure 3B-D). These data show that any interpretation of the 1-in-100 carrier rate threshold criteria limits detection of at-risk couples relative to the 176-gene panel and that the extent of reduction varies widely.

**FIGURE 3.**
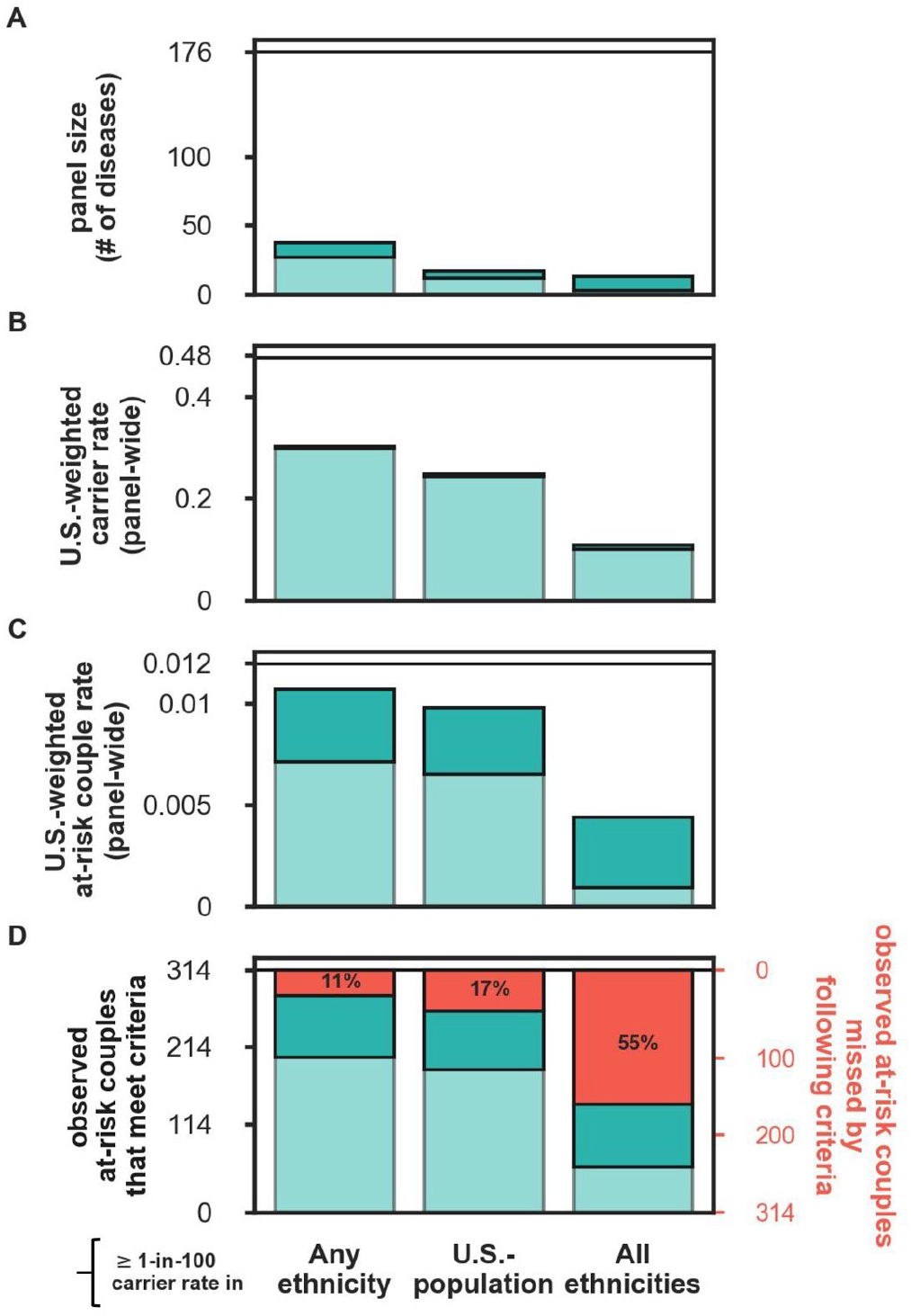
Impact of different 1-in-100 carrier-rate threshold definitions on carrier rates and at-risk couple rates. Included conditions met the first 6 ACOG criteria (Figure 1) and different definitions of the 1-in-100 carrier frequency ACOG criteria (x-axis). Shading of green bars denote different carrier rate thresholds for X-linked conditions: a carrier rate threshold of 1-in-100 (light green) and a carrier rate threshold of 1-in-10,000 (dark green). (A) The number of diseases that meet criteria. (B) U.S.-weighted panel carrier rates. (C) U.S.-weighted at-risk couple rate. (D) The number of observed at-risk couples that would be identified by the panel subset. Horizontal lines show respective numbers for the 176-condition panel.

### Panel carrier rates and at-risk couple rates saturate at large panel sizes

Because detecting at-risk couples incurs costs associated with identification of carriers, we sought to understand the quantitative interplay between the panel carrier rate and the at-risk couple rate as a function of panel size and U.S.-weighted carrier rates. The panel carrier rate increases as more conditions are added to a panel; however, this increase begins to saturate as rare conditions are added because many carriers of the rare condition were already carriers of at least one of the common conditions (Figure 4A). Growth in the panel carrier rate is rapid for panels with fewer than 18 conditions, where condition-specific carrier rates exceed 1-in-100. Even though a panel with 18 conditions has nearly 10x fewer genes than the 176-gene panel, this small panel identifies 61.0% of the carriers discovered on the large panel. Adding 73 more conditions to the panel, corresponding to a carrier rate threshold of 1-in-500, identifies 80.5% of panel carriers. The remaining 85 rare conditions that complete the 176-gene panel would increase the panel carrier rate by 19.5%.

**FIGURE 4.**
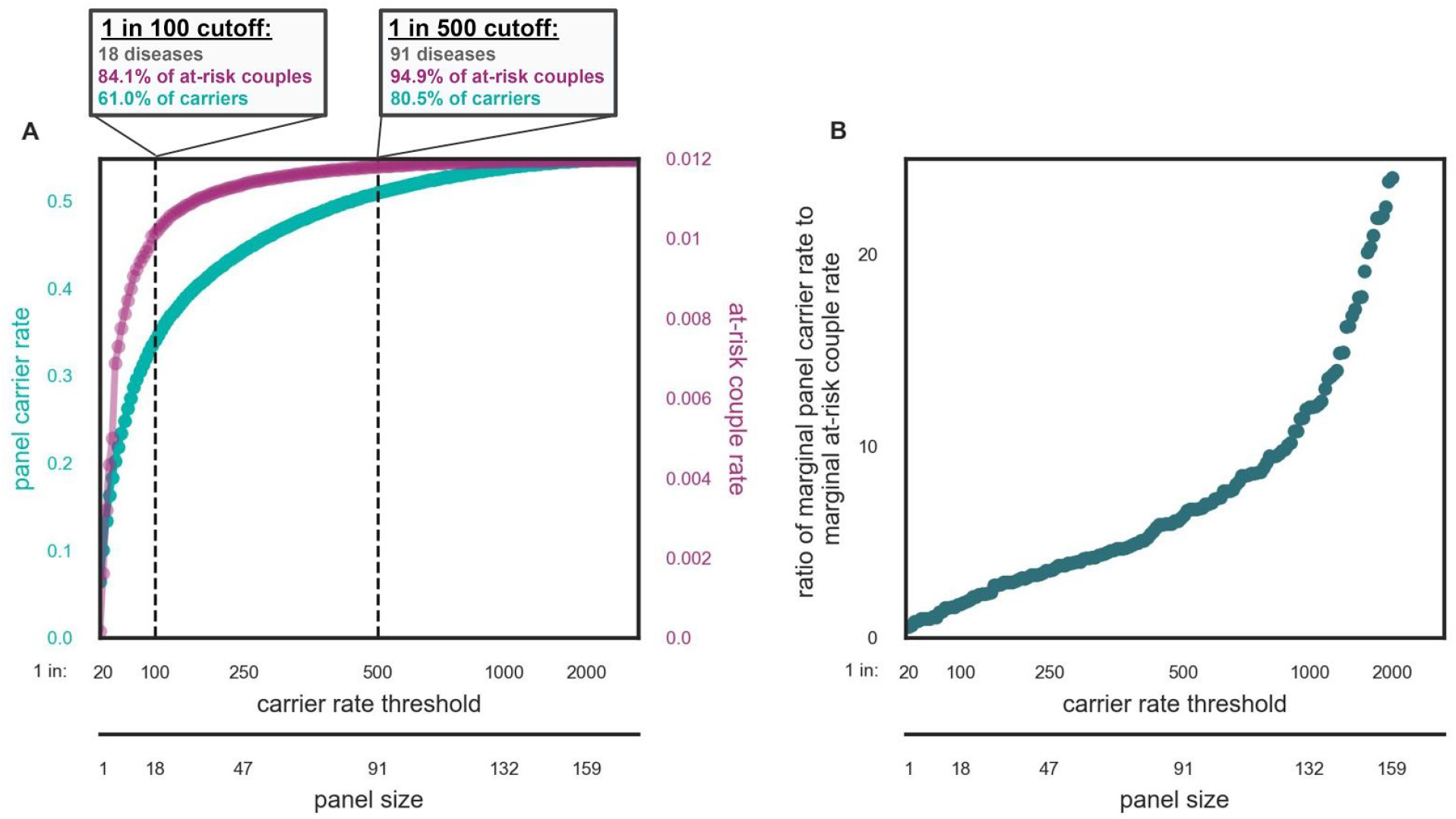
The relationship between panel size, panel carrier rate, and at-risk couple rate. (A) The panel carrier rate (green) and panel at-risk couple rate (purple) are plotted as a function of the carrier rate threshold and panel size (x-axis). Conditions are ordered from most to least prevalent based on U.S.-weighted carrier rate. Because X-linked and AR conditions are inherited differently, carrier rates for X-linked conditions were transformed to corresponding AR carrier rates based on their at-risk couple-rates. For example, for an X-linked condition with a carrier rate and at-risk couple rate of 1-in-10,000, the carrier rate would be transformed to 1-in-100. (B) Ratio of marginal panel carrier rate to marginal at-risk couple rate as a function of the panel size determined by carrier rate threshold (x-axis). The panel carrier rate was calculated from condition-specific U.S.-weighted carrier rates. At-risk couple rates were calculated as the square of the U.S.-weighted carrier rate.

Detection of at-risk couples also increases as more conditions are added to a panel and saturates as rare conditions are added to the panel (Figure 4A). However, the at-risk couple rate saturation occurs at smaller panel sizes than for the panel carrier rate because at-risk couple detection is proportional to the square of the condition carrier rate. A panel with 18 conditions accounts for 84.1% of at-risk couples. The addition of 73 conditions to the panel increases the percentage of at-risk couples identified to 94.9%, and the remaining 85 rare conditions increases the percentage of at-risk couples identified by 5.1% (Figure 4A).

We reasoned that a well-motivated carrier-rate threshold would occur when the marginal cost of screening a condition disproportionately outweighed its marginal benefit. As such, we viewed detection of at-risk couples as the benefit of carrier screening, considered identification of carriers as the cost, and quantified the relative ratio of marginal panel carrier rate to the marginal at-risk couple rate (Figure 4B). This clinically focused cost-to-benefit ratio grows as conditions become more rare, but the relationship is roughly linear down to carrier rates as low as 1-in-1000, where a subtle inflection point appears (Figure 4B). Notably, however, no conspicuous change in the ratio near a frequency of 1-in-100 is apparent.

### Estimated clinical detection rate provides an alternative for determining ECS panel rare disease exclusion criteria

Identification of at-risk couples and reduction in residual risk for patients who screen negative can only occur for a given condition if the clinical detection rate is high. Because direct measurement of clinical detection rate is challenging for rare conditions, we developed a statistical framework that estimates U.S.-weighted clinical detection rate by modeling whether a sufficient number of cases has been reported in the literature to interpret the pathogenicity of observed variants (see Methods). A schematic of the methodology is shown in Figure 5A and B.

**FIGURE 5.**
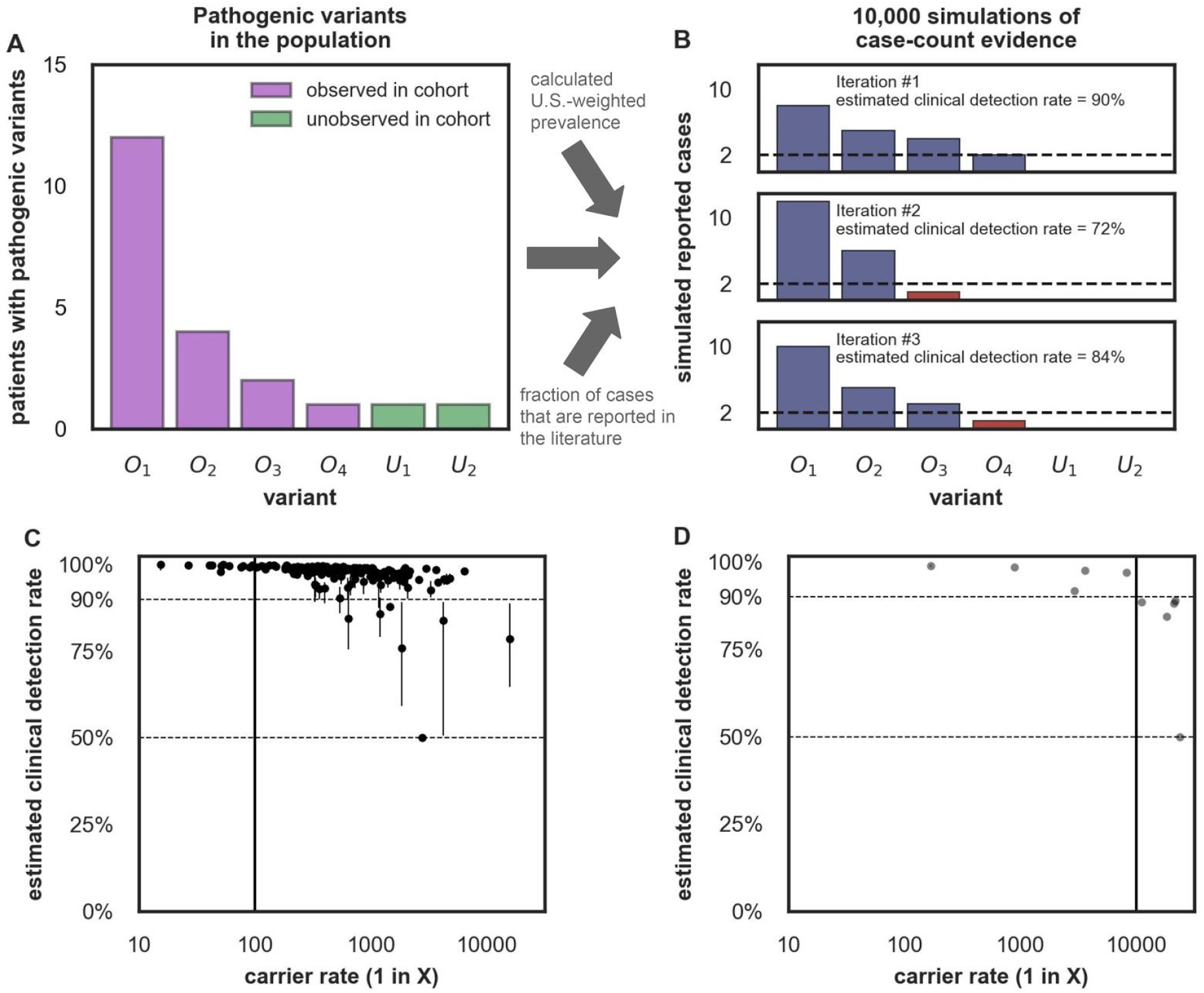
Estimation of clinical detection rate. (A-B): A model schematic showing how clinical detection rate is estimated for a hypothetical AR condition. We assume a condition has five pathogenic variants with variant frequencies shown and a carrier rate of 1-in-10,000, resulting in a prevalence of 1 in 400,000,000. (A) Assumed number of pathogenic variants, including both observed variants (purple, variants denoted with O) and a minority of unobserved variants (green, variants denoted with U). (B) Simulations of the expected number of reported cases (assuming all cases will be reported). The estimated clinical detection rate is defined as the sum of the variant frequencies for variants that can be classified as pathogenic, determined by three or more estimated case reports (shown in blue). Variants whose pathogenicity cannot be determined are shown in red or have no reported cases. (C-D) Estimated clinical detection rates for (C) AR-conditions and (D) X-linked conditions on the 176-condition panel. U.S.-weighted carrier rates and estimated clinical detection rate for each condition are shown when three reported cases are needed to determine pathogenicity for each variant. Dots show median estimated clinical detection rate and lines show corresponding 95% confidence intervals. We excluded X-linked severe combined immunodeficiency and X-linked ornithine transcarbamylase deficiency from this analysis as we have not observed any carriers of these X-linked conditions during the study period (see Supplementary Text S3). Conditions and corresponding clinical detection rate estimates are provided in Table S4.

Generally, the estimated clinical detection rate was lower for rare conditions than for common conditions, consistent with rare conditions having fewer reported cases than common conditions (Figure 5C,D). However, a rare condition with a small number of recurrent pathogenic variants may have higher estimated clinical detection rate than a more common condition with a large number of low-frequency pathogenic variants. For example, delta-sarcoglycanopathy (*SGCD*) has a carrier rate of 1-in-6,000 but only four observed pathogenic variants, and its median estimated clinical detection rate was 98%. By contrast, methylmalonic acidemia, cblA-type, (*MMAA*) has a carrier rate of 1-in-600 and 13 observed pathogenic variants, giving a median estimated clinical detection rate of 85%.

Each AR condition that had a carrier rate of 1-in-100 or greater (13 conditions) had an estimated clinical detection rate above 97% (Figure 5C). With the exception of *MMAA* described above, the remaining 123 conditions with carrier rates above 1-in-1000 had a median estimated clinical detection rate above 90% (Figure 5C); with 109 (88%) conditions having a median estimated clinical sensitivity above 97%. For the remaining 50 conditions with carrier rates below 1-in-1000, 39 conditions had median estimated clinical detection rate above 90% (Figure 5C,D), suggesting that residual risk reduction is possible for many rare conditions. Furthermore, these estimates are robust to adjustments of the assumed number of reported cases needed to interpret the pathogenicity of variants (Figures S4-S6). In sum, we demonstrate that diseases with carrier frequencies well below 1-in-100 achieve greater than 90% median estimated clinical detection rate.

## DISCUSSION

Precise and well-motivated panel content criteria are needed to ensure that ECS results maximize detection of at-risk couples and facilitate reproductive decision-making.^1,3,5^ Here, we have shown the first data-driven evaluation, to our knowledge, of the impact of ACOG guidelines^1,5^ for ECS panel content on detection of carriers and, critically, at-risk couples. Our analysis leveraged screening results from a diverse cohort of over 50,000 average-risk patients screened for 176 recessive conditions.

Quantitative inclusion criteria encourage consistency in clinical ECS offerings because numbers are unambiguous; however, the choice of which numbers to use should be transparently data-driven, such that the implications of guidelines are clear. The 1-in-100 carrier-rate threshold for condition inclusion proposed by ACOG aimed to address a trade-off between achieving high clinical utility and minimizing anxiety, but, importantly, data were not presented to support this particular threshold.^1^ Stevens et al. supported this threshold because a woman who screens positive for a recessive condition with a carrier rate of less than 1-in-100 would have a reproductive risk of less than 1-in-400, similar to risk cutoffs for common prenatal screening tests such as maternal serum screening.^8^ However, their justification of this threshold is unfounded because it is based on an unintended use of ECS (the testing of a single patient) rather than the intended use (the sequential testing of a couple, supported by medical guidelines^1,18^). When used as intended, a woman who screens positive for a condition with carrier rate of 1-in-100 would likely not need to proceed immediately to diagnostic testing because she can refine her residual risk through testing of her partner: if he screens positive, the reproductive risk is 1-in-4; if he screens negative, the risk can be as low as 1-in-250,000 (depending on carrier frequency and detection rates).^19^

We directly evaluated the clinical impact of the 1-in-100 carrier-rate threshold for inclusion of conditions on ECS panels and found that it warrants revisiting. The criterion does not specify in which ethnicities the 1-in-100 threshold should be satisfied, and does not offer guidance for X-linked conditions, which contribute disproportionately to at-risk couple detection compared to AR conditions with similar carrier rates. Critically, our analysis demonstrated that any definition of the 1-in-100 carrier rate threshold limits detection of at-risk couples.

We introduced the concept of a “panel carrier rate”—defined as the proportion of patients who were carriers of at least one condition on the panel—because it describes how frequently single-gene partner testing is warranted in a sequential-screening workflow and, thus, reflects the logistical and economic costs incurred to reap the clinical benefit of identifying at-risk couples. Our analysis showed that a 1-in-100 carrier-frequency threshold was not a conspicuously clear choice based on the data: both the panel carrier rate and at-risk couple rate saturate as rare diseases are included in ECS panels (Figure 4A), yet this saturation occurs beyond the 1-in-100 carrier-rate threshold for both panel carrier rates and at-risk couple rates. We additionally saw no clear point in which the marginal cost of screening and detecting carriers far exceeded the marginal benefit of detecting at-risk couples (Figure 4B). Therefore, the clinical burden associated with identification of carriers and testing of their partners is not substantially reduced by excluding rare conditions from an ECS panel. Taken together, these results show that the 1-in-100 carrier rate threshold will limit detection of at-risk couples without substantially reducing clinical burden.

Despite the drawbacks of the 1-in-100 criterion, it is important to determine when a condition is too rare to include on an ECS panel. For instance, if a condition is so rare that the pathogenicity of variants cannot be interpreted, then the test will have a 0% clinical detection rate, rendering screening useless. All conditions on an ECS panel should have a high analytical detection rate and clinical detection rate that together minimize residual risk.^3,8^ Since the analytical detection rate is >99.9% for most conditions,^13^ we suggest that ECS panel-content criteria should focus on defining an acceptable clinical detection rate. Whereas a carrier-rate threshold alone is problematic because no direct relationship exists between carrier rate and residual risk, a clinical detection rate is a direct measure of variant interpretability and an indirect measure of disease prevalence. We implemented a statistical method to estimate clinical detection rates for conditions on the 176-condition panel and demonstrated that conditions with carrier rates as low as 1-in-1000 have a greater than 84% estimated clinical detection rate (Figure 5). Clinical detection rates do fall as conditions become less common; thus, it is incumbent upon laboratories offering large ECS panels to demonstrate the clinical detection rate of screened conditions.

Many factors influence ECS panel content, and our study has been purposefully limited to an evaluation of clinically useful metrics. Other factors that could affect panel size include the clinical utility of screened diseases and the economic feasibility of testing a large panel. However, we have recently demonstrated that disease severity, not rarity, is a driver of ECS clinical utility,^20^ and that the high-throughput of NGS testing enables inclusion of additional conditions at a low cost.^21^ Additional limitations include that we did not explicitly evaluate the increased clinical burden associated with screening rare conditions including partner testing, genetic counseling, and patient anxiety.^2^ Further, although our estimation of clinical detection rate attempted to account for unobserved pathogenic variants, we made assumptions about the number and frequency of unknown variants, and future research is needed to refine these estimates.

In summary, we have shown the first data-driven evaluation in a large patient cohort of the impact on carrier and at-risk couple detection of ECS panel condition-inclusion criteria recommended by medical societies. While guidelines are needed to ensure high clinical utility of ECS panels, we showed that the 1-in-100 carrier rate threshold is not supported by data and limits detection of at-risk couples without minimizing residual risk. Instead, we propose that the clinical detection rate of a severe condition is a better determinant of its suitability for screening than its carrier rate alone.

## ACKNOWLEDGEMENTS

We thank Katie Johansen Taber, Kenny Wong, Bryan Dechairo, and Eric Evans for comments on the manuscript, and Anthony Gregg and Michael Guo for helpful discussions.

